# Building Models of Functional Interactions Among Brain Domains that Encode Varying Information Complexity: A Schizophrenia Case Study

**DOI:** 10.1101/2020.12.10.420208

**Authors:** Ishaan Batta, Anees Abrol, Zening Fu, Adrian Preda, Theo G.M. van Erp, Vince D. Calhoun

**Affiliations:** Center for Translational Research in Neuroimaging and Data Science (TReNDS): Georgia State University, Georgia Institute of Technology, and Emory University, Atlanta, USA; Department of Psychiatry and Human Behavior, University of California Irvine, Irvine, California, USA; Dept. of Electrical and Computer Engineering, Georgia Institute of Technology, Atlanta, USA

**Keywords:** Multilayer Perceptron, Bayesian Optimization, Hyperparameter Optimization, Schizophre, nia, fMRI, Functional Connectivity, Subdomain Analysis

## Abstract

Revealing associations among various structural and functional patterns of the brain can yield highly informative results about the healthy and disordered brain. Studies using neuroimaging data have more recently begun to utilize the information within as well as across various functional and anatomical domains (i.e., groups of brain networks). However, most whole-brain approaches assume similar complexity of interactions throughout the brain. Here we investigate the hypothesis that interactions between brain networks capture varying amounts of complexity, and that we can better capture this information by varying the complexity of the model subspace structure based on available training data. To do this, we employ a Bayesian optimization-based framework known as the Tree Parzen Estimator (TPE) to identify, exploit and analyze patterns of variation in the information encoded by temporal information extracted from functional magnetic resonance imaging (fMRI) subdomains of the brain. Using a repeated cross-validation procedure on a schizophrenia classification task, we demonstrate evidence that interactions between specific functional subdomains are better characterized by more sophisticated model architectures compared to less complicated ones required by the others for optimally contributing towards classification and understanding the brain’s functional interactions. We show that functional subdomains known to be involved in schizophrenia require more complex architectures to optimally unravel discriminatory information about the disorder. Our study points to the need for adaptive, hierarchical learning frameworks that cater differently to the features from different subdomains, not only for a better prediction but also for enabling the identification of features predicting the outcome of interest.

## 1 INTRODUCTION

Numerous works have studied brain disorders by employing multilayered machine learning (ML) approaches to neuroimaging data. In most cases, the main focus of these studies is to increase the accuracy with which the subjects having a certain condition can be classified from unaffected controls. However, not many studies focus on probing the predictive power of features encoded by the neuroimaging data. In addition to improving the classification capabilities of developed classifiers, it is equally important to localize the brain regions or groups of brain regions (subdomains) that are the most discriminative for a given disorder. Recently, deep learning classifiers have been applied widely in studies involving the use of neuroimaging data for classification. So far, previous work has used the desired features directly as a single input to the learning framework without any subdomaining of the feature set (Srinivasagopalan et al., 2019; Ulloa et al., 2015; Han et al., 2017). The prevailing methodologies inherently employ two implicit assumptions which may not necessarily hold, namely a homogeneity in the feature set for use as a single input set and a high uniformity in the complexity of feature interactions between subdomains, ignoring the need for flexible architectures. Mainstream architectures based upon these assumptions leave little room for interpretation because the parameters learned in multi-layered learning models are non-linear combinations of the input features. While using branched architectures can be of help with interpretation in terms of subdomains in the data, various studies which use multi-branched architectures on neuroimaging data have mainly focused only on accuracy improvement rather than interpretability and used these architectures to cater to the multimodal (Ulloa et al., 2015, 2018) and even multi-atlas scenarios (Zeng et al., 2018). Even in most of these cases, there is little flexibility in the architectures in terms of the depths of the branches. Introducing such flexibility in model complexity of multi-branched architectures allows for studying subdomains in terms of the nature of information they encode towards discriminating between two or more groups of subjects. No studies have explored the use of architectures designed to treat subdomains in the data differently and also reflect the variability with which different subdomains in the data encode predictive information. To overcome these limitations, this study introduces flexible ML architectures that take into account the variations in the complexity of interactions between subdomains in the data. Our approach demonstrates the need for as well as the benefit of disengaging the implicit assumption of feature homogeneity and uniform complexity in the nature of predictive information.

It is vital to identify interactions of functional and anatomical networks of high predictive value towards certain targeted applications. Moreover, it is equally important to study the diversity in the way this predictive information is encoded in the data with respect to the anatomical and functional subdomains of the brain. As mentioned before, the latter variation necessitates the use of architectures that are capable of incorporating inputs with varying information complexities in the data subdomains. Given the complex manner in which interactions between various regions of the brain occur, it is reasonable that certain subdomains of the brain may need deeper architectures for better prediction, which may be indicative of a greater degree of non-linear interactions in these subdomains. In this study, we analyze the pattern in which various subdomains for functional magnetic resonance imaging (fMRI) data encode information in a schizophrenia classification task. We use a multilayer perceptron (MLP) classifier for studying the functional connectivity features associated with schizophrenia classification. Towards this goal, we use the intra-network and inter-network connections at the level of subdomains, termed as subdomain interactions (SDIs) in this paper to create separate input layers in the multi-branched architecture. The input layers in our multi-branched architecture are followed by a variable number of hidden layers in each of the branches before a late fusion step as detailed in the methods section. By optimizing over this flexible multi-branched MLP architecture search space, we show that different subdomain interactions encode discriminatory information with variable complexity and certain subdomain interactions associated with schizophrenia consistently need more complex frameworks while others require simpler ones.

One of the potential concerns with allowing such flexibility in the depth is the vast architecture search space generated due to the multiple parameters involved. Performing optimization over this space is a hyperparameter optimization problem, a well studied field in machine learning (Shahriari et al., 2015; Luo, 2016; Bergstra et al., 2013b, 2011). It is computationally impractical to linearly traverse through exponentially huge hyper-parameter search spaces with standard methods such as random search or grid search. Bayesian optimization frameworks resolve this issue by heuristically traversing parts of the search space that are more likely to cover a solution close to the optimal one. Here we employ a Bayesian method known as the Tree Parzen Estimator (TPE) to realize the hyper-parameter optimization stage (Bergstra et al., 2011). Starting with a set of initial randomly chosen points in the search space, the TPE algorithm traverses new points in each of its iterations using a simpler (i.e., faster) surrogate function and calculating a metric for the expected improvement in classification accuracy subsection 2.4. In this way, the search space is traversed without having to compute the actual function (in this case, the validation accuracy of the architecture) and selecting a new point which would be more optimal with high probability. Analyzing the final architecture returned by the TPE procedure can reveal specific associative patterns corresponding to each subdomain. By studying the patterns in the associations of subdomains in the optimized architectures, we illustrate that allowing for and optimizing over subdomain specific variation in architectures not only enables superior prediction, but also reveals how certain subdomain interactions bear higher complexity of information while others have less complex information. Moreover, with the rapid increase in multimodal datasets for mental disorders, it is crucial to develop methodologies synthesizing features from subdomains spanning more than one modality. Our study also gives initial insights towards the need for developing such flexible frameworks for multimodal studies. Frameworks catering to and analyzing subdomain variation can be of immense use in various fields of study.

The flow of the paper, shown in Figure 1 and detailed in methods section, is as follows: a) Spatially constrained independent component analysis (scICA) on the preprocessed fMRI dataset is used to calculate the functional connectivity for pairs of components of interest subject-by-subject; b) Categorizing the feature set of component-component functional connectivity into subsets based on subdomains (brain networks) of the two components involved (the inter-network or inter-network connections are termed as subdomain interactions (SDIs) throughout the paper); c) Using the subdomained features as inputs to the multi-branch MLP architecture with flexible depth, with hyper-parameter optimization (TPE algorithm) on the architecture search space to determine the optimal variability in depth for each SDI (Figure 1b); and d) Comparison of the performance of TPE with baseline methods and analysis of patterns associated with certain SDIs in the validated optimal architectures returned by the TPE procedure. We demonstrate the working, performance and interpretation of the TPE algorithm in the results section. Lastly, we discuss the interpretation of these results in the discussion section.

**Figure 1.**
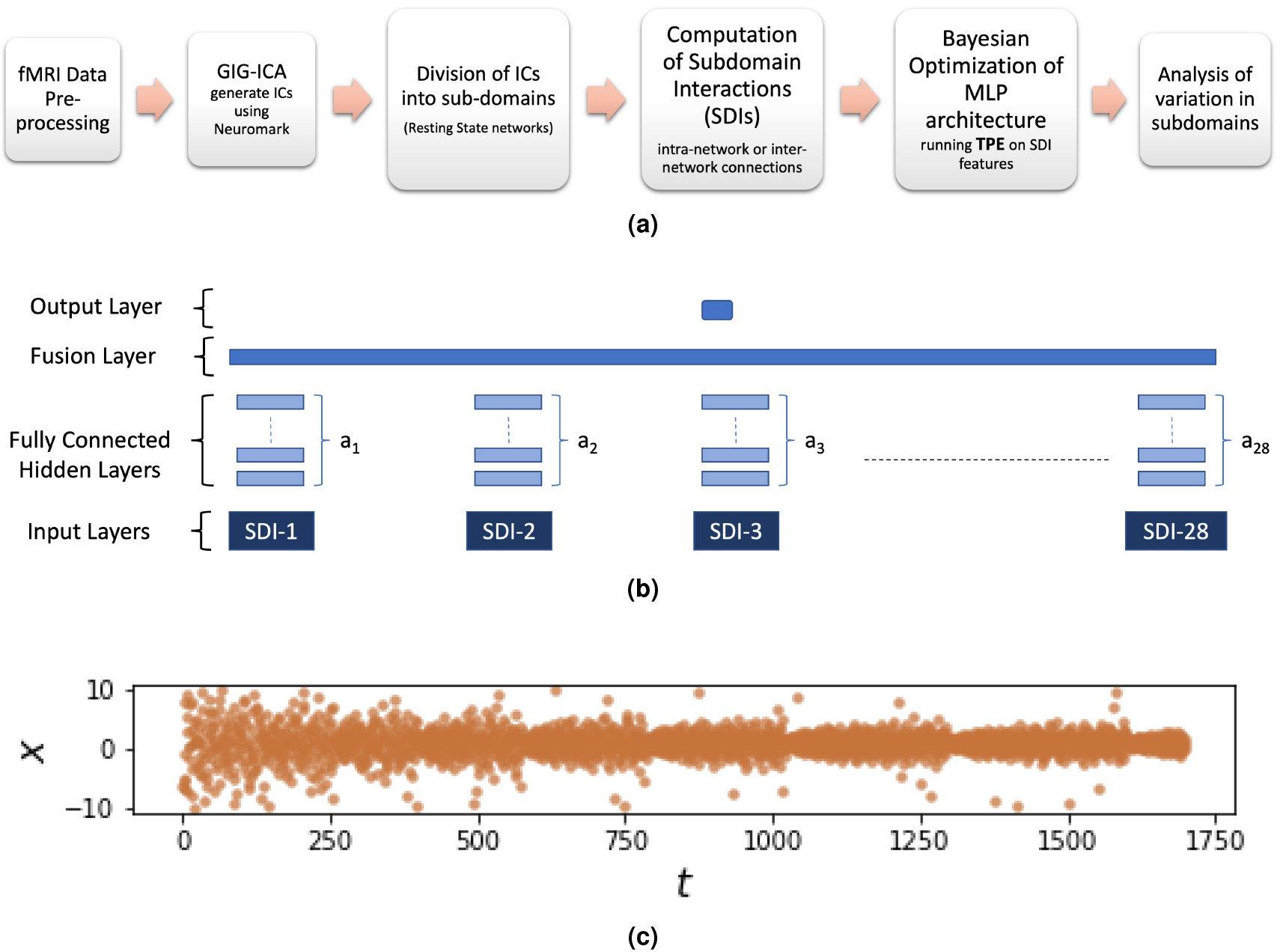
(a) A step-by-step description of the whole analysis. (b) An architecture in the search space that TPE optimizes over, defined by the vector 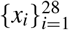, with *x_i_* ∈ {0,1,2} representing the number of fully connected hidden layers on top of the input node corresponding to data from the i-th subdomain interaction (SDI). Data for each SDI is a sub-matrix of the full static functional connectivity matrix with connections from participating subdomain(s). (c) TPE search space traversal on a toy example with a quadratic cost function (*x* − 1)^2^ to narrow down the search closer to the optimal value.

## 2 METHODS AND MATERIALS

### 2.1 Datasets and Pre-Processing

This study uses two independent datasets, the Function Biomedical Informatics Research Network (fBIRN) dataset (Keator et al., 2016) and the Center of Biomedical Research Excellence (COBRE) dataset (Aine et al., 2017). Subjects with large head motion (≥ 3 deg and ≥ 3mm) during the scan and with functional data leading to bad full brain normalization were excluded. After applying this criteria for exclusion, the fBIRN dataset consisted of 160 healthy controls (HC) with age 19 – 59 years (mean 37.04 ±10.86), 45/115 females/males and 151 subjects who had schizophrenia (SZ) with age 18 – 62 years (mean 38.77 ±11.63), 36/115 females/males, whereas the COBRE dataset consisted of 89 healthy controls (HC) with age 18 – 65 years (mean 38.09 ±11.67), 25/64 females/males and 68 SZ with age 19 – 65 years (mean 37.79 ±14.45), 11/57 females/males. For avoiding confounding effects, HC and SZ subjects of both datasets were matched by age and gender (age: *p* = 0.1758 (fBIRN), 0.8874 (COBRE); gender: *p* = 0.3912 (fBIRN), 0.0794 (COBRE)).

Data collection for fBIRN dataset was done using 3-T Siemens Tim Trio scanners for six out of seven sites and 3-T General Electric Discovery MR750 scanner for one site. The same resting-state parameters were used across the scanners a standard gradient echo-planar imaging (EPI) sequence, repetition time (TR)/echo time (TE) = 2000/30 ms, voxel spacing size = 3.4375 × 3.4375 × 4 mm, slice gap = 1 mm, flip angle (FA) = 77°, field of view (FOV) = 220 × 220 mm, number of excitations (NEX) = 1, and number of volumes = 162. Participants had their eyes closed and were instructed to rest quietly during the scanner. The COBRE data was collected on a single site using a 3-T Siemens Trio scanner. For the COBRE data, a gradient-echo EPI sequence was used to acquire T2-weighted functional images with the following parameters: TE =29 ms, TR = 2000 ms, flip angle (FA) = 75°, slice thickness = 3.5 mm, slice gap = 1.05 mm, field of view 240 mm, matrix size = 64 × 64, voxel size = 3.75 mm × 3.75 mm × 4.55 mm and number of volumes = 149. For the duration of the scan, subjects were instructed to keep their eyes open and passively stare at a central cross.

For both the datasets, the statistical parametric mapping (SPM12, http://www.fil.ion.ucl.ac.uk/spm/) toolbox based on the Matlab 2016 platform was used to preprocess fMRI data. To ensure equilibrium of the signal and adaptation of the subjects to scanner noise, the first five scans were removed. The SPM toolbox was used for performing slice timing correction and rigid body head motion correction. Warping of the fMRI data into the standard Montreal Neurological Institute (MNI) space was done using an echo-planar imaging (EPI) template. The data were slightly resampled to 3 × 3 ×3 mm^3^ isotropic voxels. For smoothing the data, a Gaussian kernel with a full width at half maximum (FWHM) of 6 mm was used. This was followed by further feature extraction detailed in subsequent sections.

### 2.2 Connectivity Features and subdomain Interactions

To estimate intrinsic connectivity networks from the fMRI data, the scICA approach described in (Du and Fan, 2013) with the Neuromark (Du et al., 2019; Y. Du and Calhoun, 2020) template as reference maps was used. ICA was run for a model order of 100, out of which 53 consistent and reproducible independent non-artifactual components (ICs) were retained. The functional connectivity matrix was computed based on the static correlation between the time courses of pairs of two ICs. All the 53 ICs have been arranged into 7 distinct functional subdomains (disjoint exhaustive sets of ICs), namely the areas falling under the following subdomains: default mode network (DMN), visual (VIS), auditory (AU), cognitive control (CC), sensorimotor (SM), cerebellar (CB) and sub-cortical (SC). Based on this categorization, 28 subdomain interaction (SDI) features were created where each SDI represents the set of intra-network or inter-network connections between all possible pairs of ICs for a given pair functional subdomains or pair of functional subdomains respectively (See Figure 2). As an example, the features in the SDI named DMN-VIS refer to entries from the functional connectivity matrix (fMRI time-series correlations) corresponding to the connections between the DMN and VIS subdomain scICA components. Such connections are termed as inter-network connections. On the other hand, the intra-network connections comprise connections within the same domain, for example, the SDI named DMN-DMN would correspond to the connections where both of the scICA components belong to the DMN subdomain. Thus, connectivity features for the 28 SDIs consist of 21 sub-matrices for the inter-network connections and 7 sub-matrices for intra-network connections. Hence the SDI features are simply sub-matrices of the 53 × 53 functional connectivity matrix created using 53 scICA components.

**Figure 2.**
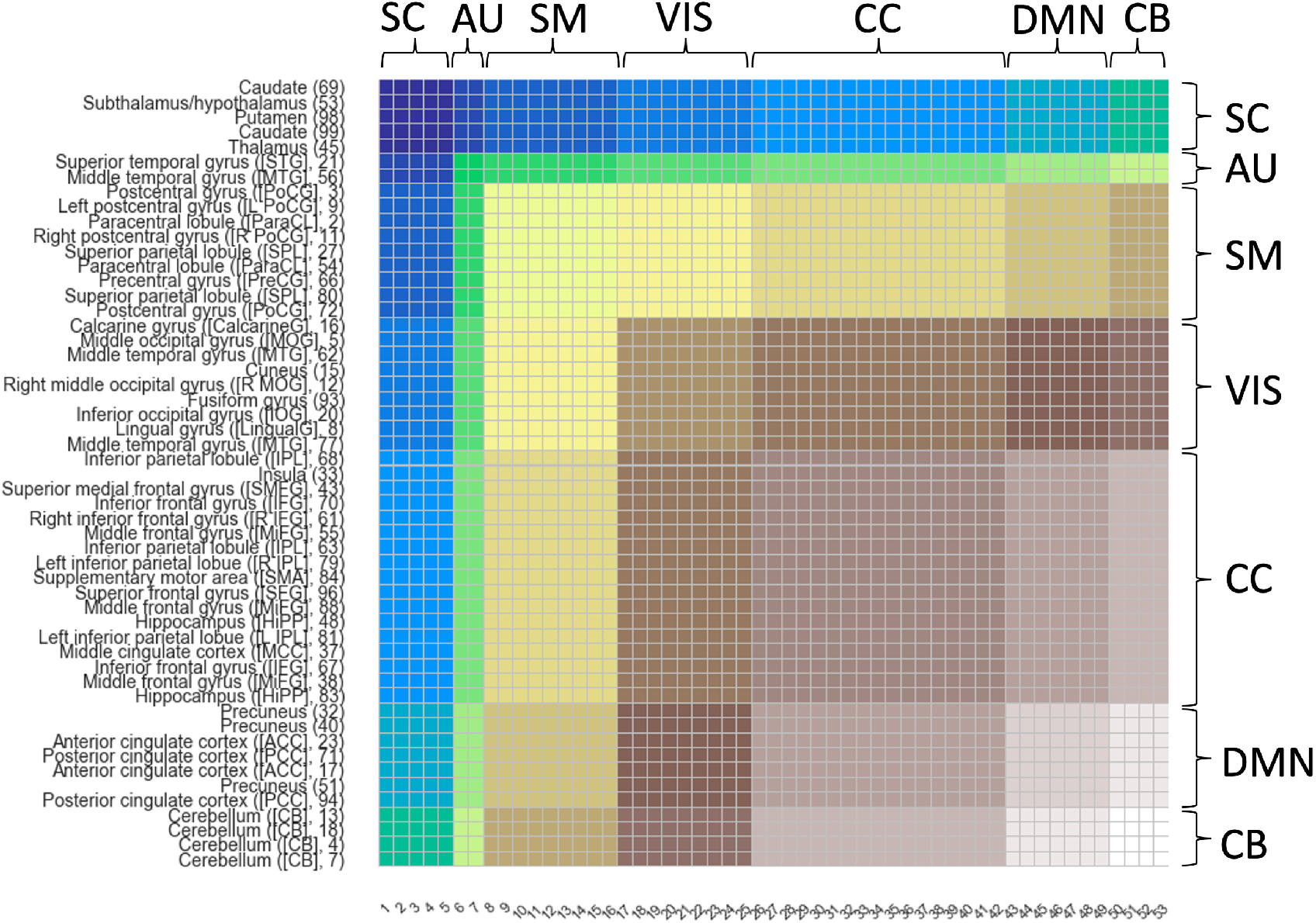
Illustration of how the functional connectivity matrix for the 53 components (53 ×53 in size) is divided into subdomain interactions (SDIs) that form the input to the branched Multilayer Perceptron (MLP) architecture. Each colored sub-matrix represents a certain SDI i.e., the set of functional connectivity values between components of a given pair of subdomains. The 7 subdomains include: default mode network (DMN), visual (VIS), auditory (AU), cognitive control (CC), sensorimotor (SM), cerebellar (CB) and sub-cortical (SC). A total of 28 SDIs, which are sub-matrices of the full functional connectivity matrix, are colored in different colors. Since the number of subdomains is 7, a total of ^7^C_2_=21 Out of the 28 SDIs correspond to inter-network connections (e.g. DMN-VIS) while 7 correspond to intra-network connections (e.g. DMN-DMN).

### 2.3 Architecture Search Space

A multilayer perceptron (MLP) architecture with multiple branches was used with input layer consisting of the 28 SDI features from the 7 subdomains (sets of ICs) as described in subsection 2.2. At the input level, the architecture has 28 branches for each SDI i.e., a sub-matrix of the 53×53 static functional connectivity matrix. Each input layer is then fed into a variable number of fully connected hidden layers and a late fusion layer, finally followed by a fully connected layer (Figure 1b). In each branch, a constant factor of 0.1 was used to reduce the number of units in subsequent hidden layers, starting with the input layer. Optimizing the variation in the number of fully connected layers in the branch corresponding to each SDI can give insights into understanding the nature of information that is encoded by the component interactions for the subdomains involved in the SDI. SDIs which consistently require no or a small number of layers in an optimized architecture can be said to contain more linear or direct information for the task at hand (schizophrenia classification), whereas the SDIs which require a higher number of fully connected layers can be said to contain more complex and indirect information. For controlling the model complexity, the number of fully connected hidden layers in each branch were limited to vary between 0 to 2. In the exponential search space thus generated with a total of 3^28^ possible configurations, a 28 length vector **x** can be used to represent each point, with each element *x_i_* ∈ {0, 1, 2} denoting the number of fully connected layers in the branch for the *i*-th SDI (Figure 1b).

### 2.4 Tree-structured Parzen Estimator (TPE) for Hyper-Parameter Optimization

Optimizing hyper-parameters that span an exponential search space has been a well-studied problem in machine learning. In similar lines, Bergstra et al. (2011) had proposed the TPE algorithm which is used for hyper-parameter optimization in this work. TPE is a sequential model-based optimization (SMBO) framework which essentially estimates the conditional and marginal probabilities *p*(*x|y*) and *p*(*y*) to perform hyper-parameter optimization on hyper-parameters *x* and cost function *y*. After spanning an initial set of points selected randomly from the search space and computing the cost function for them, TPE utilizes this information to create a surrogate cost function. The surrogate function is then used to select the next set of points in the traversal sequence based on the estimates of the cost function. Towards this goal, two groups made up of the upper quartile and the lower quartile are created based on a splitting value *y*\* of the cost function at the randomly selected points. The two probability density functions, *g*(*x*) and *l*(*x*) for the upper and lower quartiles of the cost function respectively is then defined as in Equation 1:

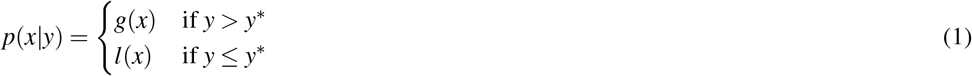

Having estimated *l*(*x*) and *g*(*x*), the subsequent iterations of the TPE algorithm involve the optimization of the Expected Improvement function defined in Equation 2:

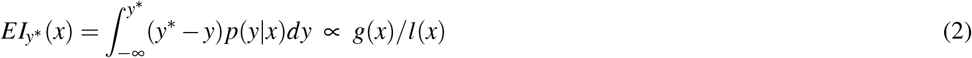

The expected improvement function *EI_y*_*(*x*) can be shown to be proportional to *l*(*x*)/*g*(*x*) (Bergstra et al., 2011). This means that if points are sampled from *l*(*x*) with high probability and from *g*(*x*) with lower probability, then the expected improvement is maximized, thus minimizing the cost function. In each iteration, the algorithm returns the point *x*\* having the largest value of *EI_y*_*(*x*) and the distributions *l*(*x*),*g*(*x*) are updated accordingly. An example showing how TPE works is provided in Figure 1c and indicates convergence towards the true optimal on a quadratic cost function (*x* − 1)^2^ optimized for a real valued parameter *x*.

### 2.5 Analysis of Learned Models

Using features from each SDI as inputs, the MLP architecture with 28 branches as described in subsection 2.3 was optimized using the TPE implementation in the HyperOpt python library (Bergstra et al., 2013a). We used a repeated random sub-sampling cross-validation procedure for running the TPE algorithm for 50 repetitions on the functional connectivity data, resulting in a set of optimized architecture vectors 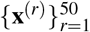. The mean across repetitions of the validation accuracy has been plotted against TPE iterations in Figure 4.

The final prediction model, represented by the vector **x**\*, was created using the most frequent number of hidden layers across repetitions for each SDI in the TPE-optimized architectures. The final architecture vector **x**\* is defined as, 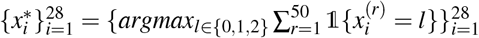, where **x**^(*r*)^ represents the TPE-optimized architecture for the *r*^th^ repetition and 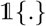 is the indicator function. The multi-branched MLP architecture generated using the final architecture vector **x**\* was run on a held-out test data for 50 repetitions, resulting in 50 test accuracy values.

### 2.6 Comparison with Baseline Methods

To study the performance of the TPE-optimized architecture, a comparison was done with baseline methods run for 50 repetitions on held-out test data. These methods included simple machine learning models like logistic regression (LOG), support vector machine (SVM) and random forest classifier (RFC). For an even closer comparison, we also used three types of multilayer perceptron (MLP) architectures as baseline. These included two multi-branched MLP architectures (UNIF0 and UNIF1) but with uniform depth for each branch and also a simple MLP architecture (MLP1) with a fully connected layer but without any branches. Note that the MLP1 architecture has all the static functional connectivity features in the input layer, which are further fed to a fully connected hidden layer without any branching. While the MLP1 architecture serves as a baseline to compare multi-branched vs. non-branched architecture scenarios, the UNIF1 and UNIF0 architectures serve as a baseline for comparing uniform vs. flexible depth in the branch corresponding to each SDI. Thus, against the architectures **x**^(*r*)^’s returned by TPE have variable values of layer-depth, the UNIF1 architecture essentially is an architecture in the same architecture search space of TPE algorithm, but with a constant layer-depth of 1 on top of each SDI branch before the late-fusion step. Therefore, the depth-vector for UNIF1 is 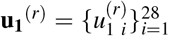 with 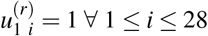, where 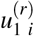 represents the number of layers used on top of *i^th^* SDI in the architecture UNIF1 for the *r^th^* repetition. Similarly, the architecture UNIF0 is designed to have no layer on top of each SDI branch before the late fusion step in the multi-branch MLP architecture i.e., the UNIF0 architecture’s depth-vector is 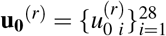 where 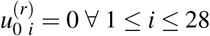.

## 3 RESULTS

### 3.1 scICA components

After running the scICA framework on the fMRI data, 53 non-artifactual, reproducible independent components (ICs) were obtained. Table 1 shows the peak coordinates for the components along with the brain region corresponding to the component. Following this, static correlations between the time courses of these 53 components were computed resulting in a 53×53 functional connectivity matrix. The 53 ICs were assigned to 7 functional subdomains or sets of components (Table 1, Figure 3). 28 pair-wise subdomain interactions (SDIs) resulting from the 7 subdomains were constructed. The SDIs are defined by the sets of intra-network or inter-network connections between all possible pairs of subdomains; thus, there were 21 SDIs corresponding to inter-network connections and 7 for the intra-network connections (Figure 2).

**Table 1.**
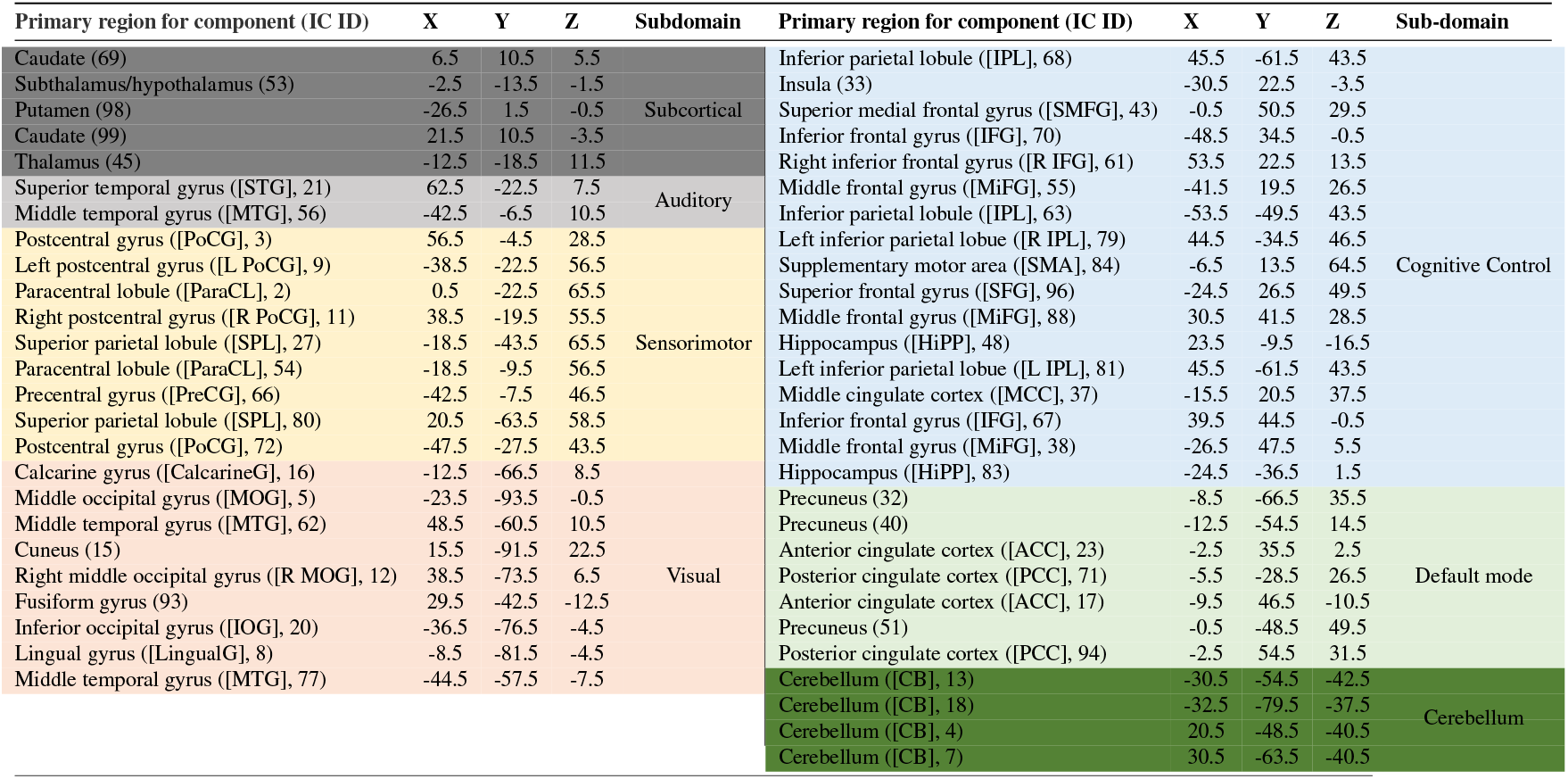
Peak coordinates and primary brain regions for the 53 components (ICs) obtained using scICA (Neuromark framework) on the fMRI time series data. The 53 components were divided into 7 subdomains (resting-state networks) as shown in the table. The time-series from every possible pair of these 53 components was used to compute the static functional connectivity (SFNC) features using Pearson correlation. The SFNC features were then divided into 28 subdomain interactions (SDIs) based on the subdomain(s) to which a given pair of components for the SFNC feature belongs (See Figure 3).

**Figure 3.**
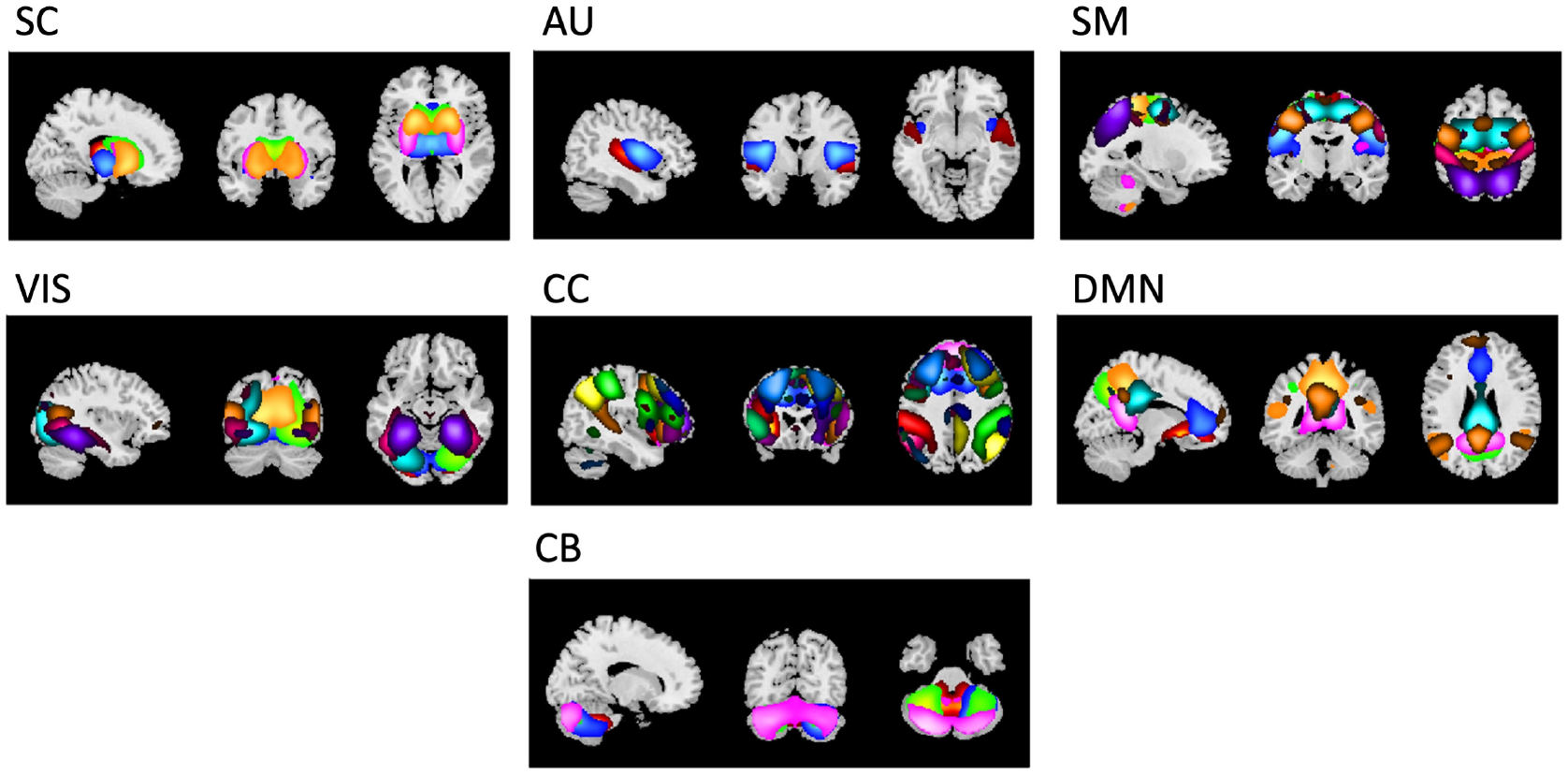
The components obtained from the scICA procedure Neuromark are shown in different colors. Each map shows the components belonging to a particular subdomains i.e., networks of brain. The 7 subdomains include: default mode network (DMN), visual (VIS), auditory (AU), cognitive control (CC), sensorimotor (SM), cerebellar (CB) and sub-cortical (SC). subdomain interaction (SDI) features built using these subdomains and components were used as input to the multi-branch MLP architecture optimized for variable branch-depth by the TPE algorithm. The SDI features are comprised of functional connectivity (static time-series correlation) between pairs of components belonging the same subdomain (intra-network connections) or different subdomains (inter-network conenctions). The 7 subdomains shown above lead to the creation of 28 SDIs, with ^7^C_2_=21 corresponding to the inter-network connections (Eg. DMN-VIS) and 7 corresponding to the intra-network connections (Eg. DMN-DMN).

**Figure 4.**
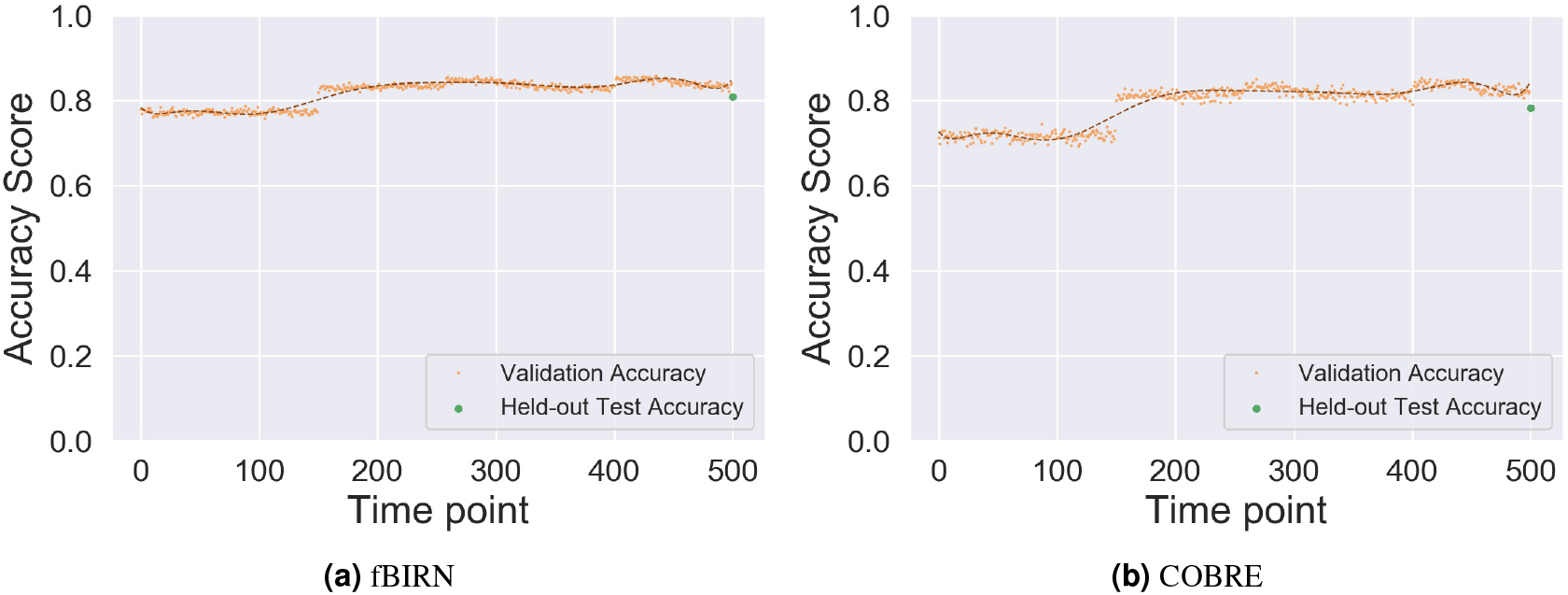
Mean validation accuracy vs. time point (iterations) for 50 repetitions of the TPE algorithm over the architecture search space depicted in Figure 1b. The mean test accuracy using the final architecture on held-out data for each repetition is also shown. The points traversed in the search space are selected based on expected improvement (Equation 2).

### 3.2 Performance using TPE

Starting with an initial set of randomly selected points in the hyper-parameter search space, the TPE optimization procedure subsequently selects new points to traverse using the expected improvement metric and the surrogate cost function which is computationally faster (subsection 2.4). Figure 4 shows the validation accuracy plotted against time (iterations). The validation accuracy, which is the objective function being optimized, increases with the TPE iterations. The procedure returns an optimal architecture for each of the 50 repetitions. The most frequently occurring number of hidden layers for each SDI across repetitions are used to define the final architecture. The test accuracy values on held-out data were obtained for 50 independent repetitions by using the final architectures constructed after running the TPE optimization algorithm. Figure 5 The final TPE models reported a slightly higher prediction accuracy on held-out test data than the baseline machine learning methods (SVM, logistic regression and random forest classifier). Specifically, we observed a mean prediction accuracy of 0.81 for fBIRN and 0.78 for COBRE for the proposed TPE-optimized architecture. For both the datasets, TPE also performs significantly better (*p* < 0.05) than the two baseline multi-branch MLP architectures with uniform depth in each branch (UNIF1 and UNIF0), while it performs slightly better than the MLP architecture without any branching (MLP1). This result shows that treating features from certain subdomains differently in terms of the complexity of architecture required can result in superior prediction performance. In fact, this improvement over the baseline methods also allows for a meaningful analysis of variability in the nature of information carried by the various subdomain interactions in the data.

**Figure 5.**
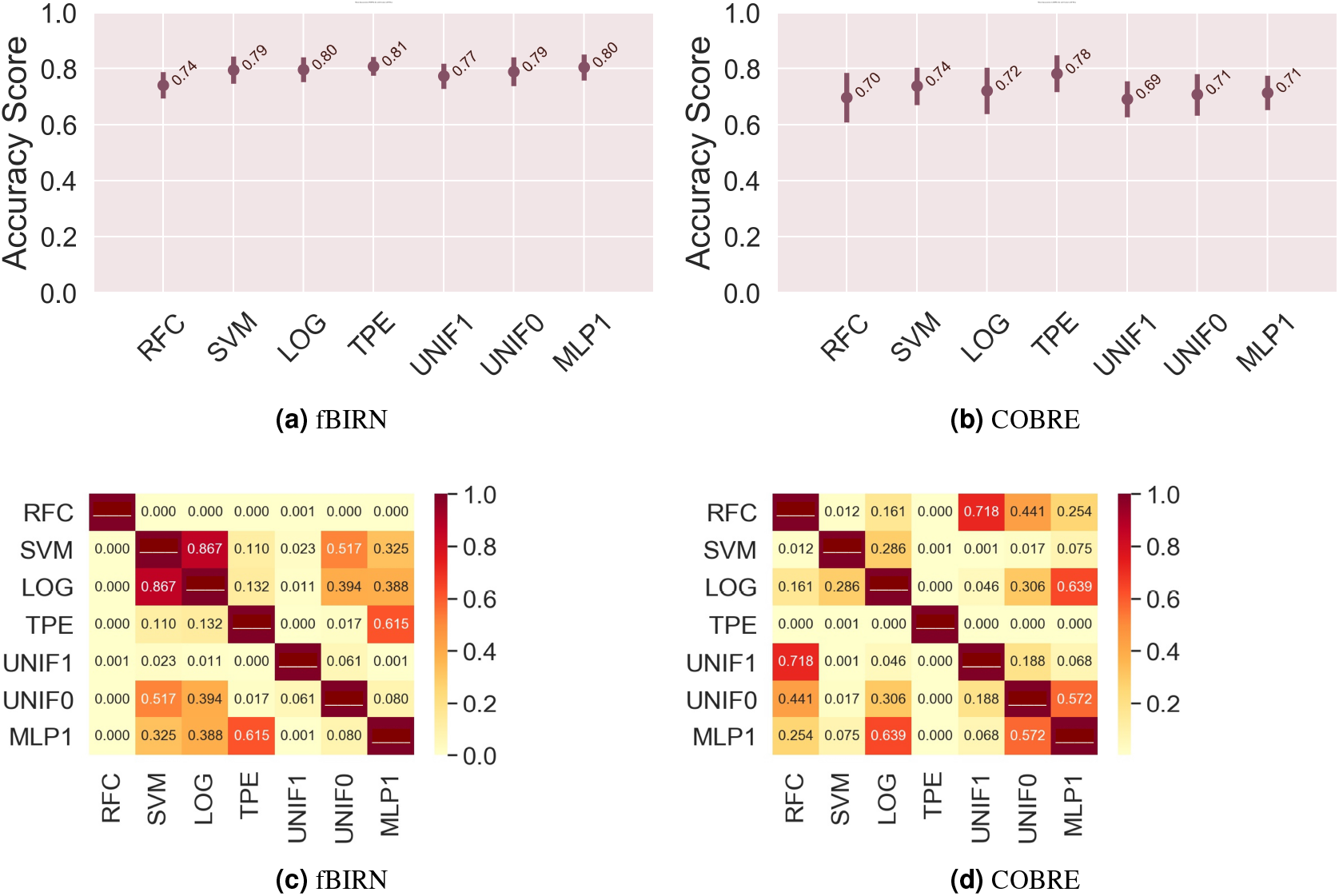
Mean validation accuracy with error-bar for 50 repetitions of the TPE-optimized final architecture in comparison to baseline methods for (a) fBIRN and (b) COBRE datasets along with p-values for two-sample t-test on the mean test accuracy across 50 repetitions, shown in (c),(d). Baseline methods include simple machine learning models (SVM, LOG, RFC) and also similar multi-branched (UNIF1, UNIF0) as well as non-branched (MLP1) MLP architectures. Simple machine learning models used were Support Vector Machine (SVM), logistic regression (LOG) and Random Forest Classifier (RFC), each optimized with grid search over parameters. Baseline MLP architectures included uniform architectures, UNIF0 and UNIF1, representing inflexible multi-branched architectures with 0 and 1 fully connected layers before the fusion step and above input layer in each SDI branch of the architecture respectively. As the third baseline MLP method, a non-branched architecture (MLP1) was used with a single fully connected hidden layer with input layer consisting of functional connectivity features (without any subdomaining). The architecture created from the repeated optimizations using the TPE procedure is termed as TPE in the plots. It can be noted that for both the datasets, the accuracy obtained by using the TPE-optimized architecture is significantly higher than the accuracies from uniformly branched architectures (UNIF0, UNIF1), indicating the need for flexible architectures. Moreover, the accuracy with TPE is slightly higher than the accuracy for the other baseline methods, showing the scope of interpretability in the optimized model in terms of certain subdomain interactions (SDIs) with higher complexity requiring deeper while others requiring shallow architectures.

### 3.3 Feature Stability

While the TPE algorithm gives the best performance, the next step would be to analyze the subdomain interactions (SDIs) requiring deeper or shallower hidden layers in the multi-branched MLP architecture. However, before checking for this characteristic variation in the SDIs, it is relevant to check whether the features learned by the optimized architecture have consistent discriminative power for each SDI in terms of their importance towards prediction. For this purpose, impurity-based feature prediction power (Louppe et al., 2013), also known as Mean Decrease Impurity (MDI), was computed by running random forest classifier on the parameters learned in the late fusion layer (Figure 1b) of the optimized TPE architecture. These were compared with the impurity-based prediction power by running random forest classifier on the features in the input layer of the network (i.e., the functional connectivity features). The comparison was done by obtaining the prediction power vectors for both the aforementioned cases and computing the cumulative prediction power for elements belonging to each subdomain interaction (SDI). The cumulative prediction power of each SDI for the prediction task, averaged over 50 repetitions of the algorithm on held-out test data is plotted in Figure 6. The results indicate that the prediction power of SDI parameters learned in the late fusion step of the final architecture is highly correlated (0.9 for fBIRN and 0.81 for COBRE) with the prediction power of the same SDI features in the functional connectivity input features. This observation suggests that the TPE-optimized architecture learned appropriate brain representations as the SDI features retained similar properties in terms of the prediction accuracy on the schizophrenia classification task as well as the importance of each SDI in the prediction. With these properties being similar, the TPE procedure that optimizes based on the depth by each SDI in the multi-branched MLP architecture, can be said to additionally provide insights into the complexity of the information stored in the SDIs. The results from the analysis of this variation in the nature of information across the SDIs (as defined in subsection 2.5) is done in the subsequent subsection.

**Figure 6.**
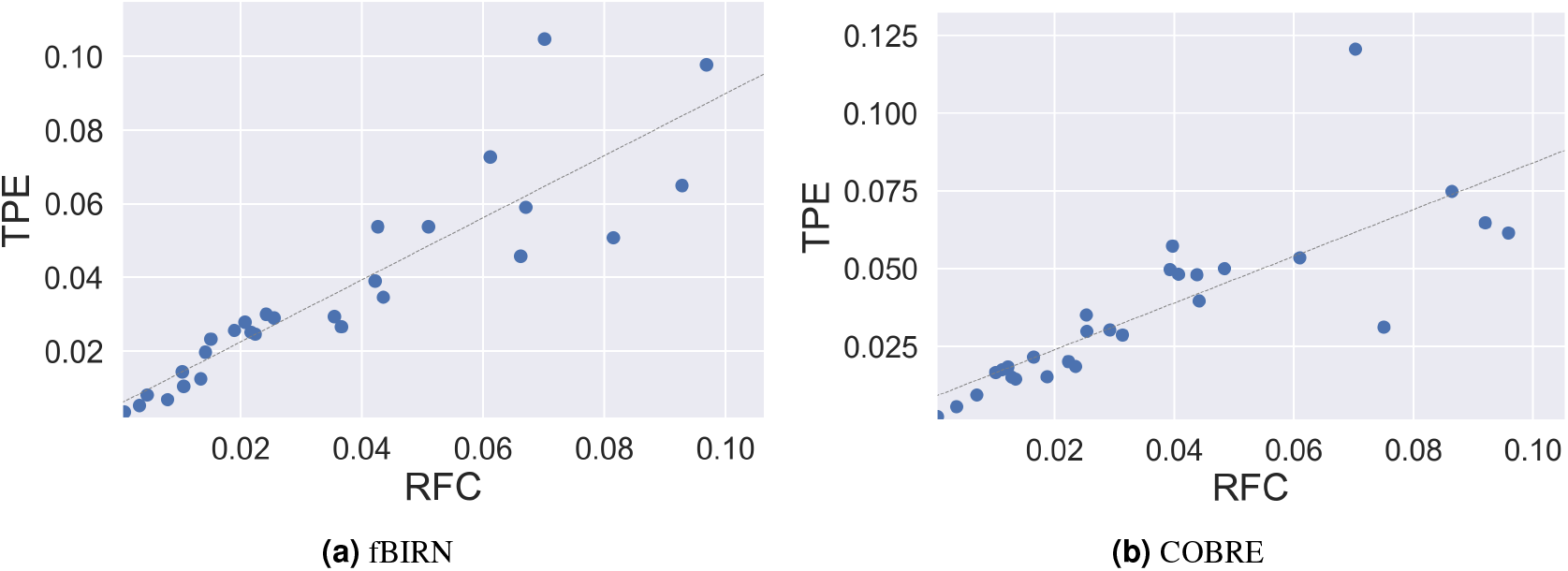
To check whether the relative importance of SDI parameters learned in the fusion layer of MLP architecture is similar to the functional connectivity features of the corresponding SDI, Random Forest classifier was used to compute purity-based feature importance on the parameters in the fusion layer as well as the input layer (connectivity features). The prediction power vector obtained for both these cases was divided into 28 bins corresponding to each SDI and summed to get the cumulative prediction power of each of the SDIs. The above plots show the cumulative prediction power of SDIs, averaged over 50 repetitions, in the learned parameters inside the fusion layer (marked as TPE on the y-axis) and in the functional connectivity features in the input layer (marked as RFC on the x-axis). There was a high correlation of 0.9 and 0.81 between these prediction power values for (a) fBIRN and (b) COBRE datasets respectively. This means that in addition to being consistent in terms of the prediction accuracy, the TPE algorithm is also consistent in terms of the importance that the SDIs have for the prediction task.

### 3.4 Analyzing Optimal Models

After having ensured that the TPE procedure for optimizing the multi-branched MLP architecture is consistent in terms of prediction accuracy on held-out test data and also shows a similar trend in the importance of SDIs, an analysis on the variation in depth required by various SDIs in the final architecture returned by TPE was done as described in subsection 2.5. This was done by considering the most frequently occurring number of fully connected layers across all repetitions of TPE for each SDI. The relationship between 7 subdomains (networks) in terms of the number of fully connected layers needed for optimal decoding of information in each pair of networks (SDI) in the final architecture is shown in Figure 7. A notable observation here is that in the final architecture of both the fBIRN and COBRE datasets, the number of fully connected layers required by each SDI is the same for 19 out of the 28 SDIs (Figure 8), which indicates a very similar common pattern across the two independent datasets in terms of complexity of information in subdomain interactions. Note that the number of such common values across two datasets can be modelled using a random variable which is distributed according to the binomial distribution *B*(*k, n, p*), where *k* is the number of successes (=19), *n* is the number of trials (=28) and *p* is the probability of a success, which in this case is the probability of having the same depth in both datasets for a particular SDI, given by 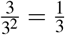. A binomial test was done to check whether the observed number of common values (=19) is significantly different from the expected number (=28/3) in the case of the binomial distribution. This test indicates that the result is significant with a *p*-value of 2.09 × 10^−4^.

**Figure 7.**
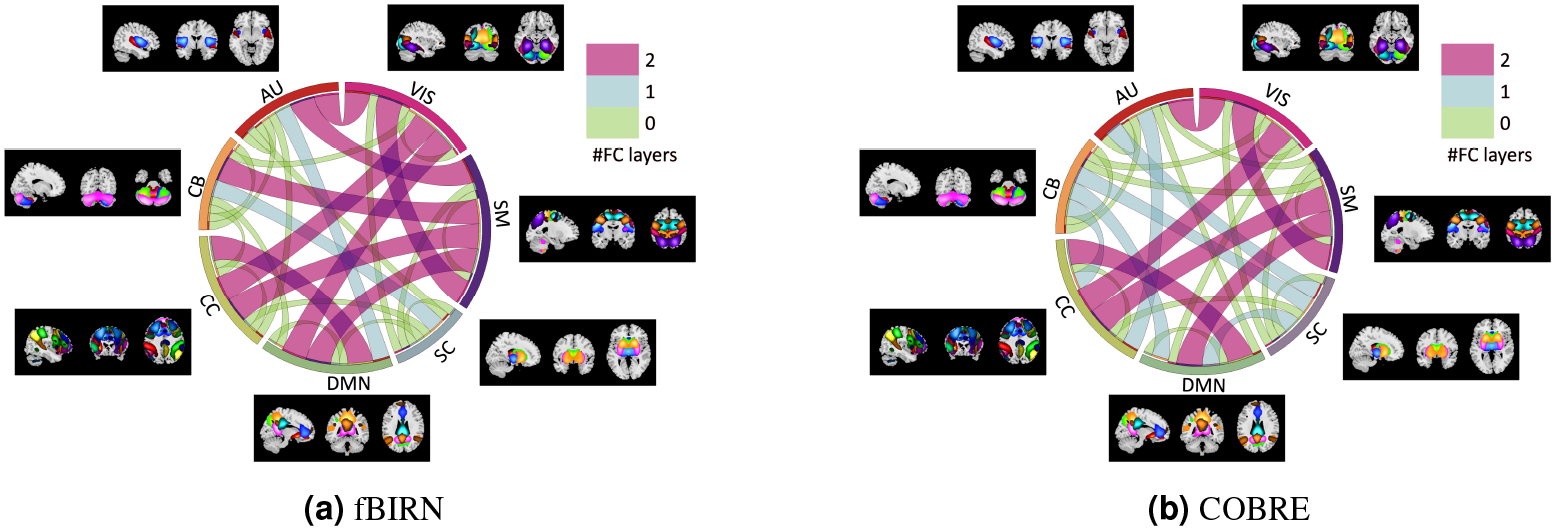
(a),(b) Visualization for the depth (in terms of the number of fully connected layers) in the final optimized architecture required by various each subdomain interactions (SDIs) (intra-network or inter-network functional connectivity). SDIs consistently requiring higher depth across repetitions of the algorithm can be said to carry more complex information towards the classification objective.

**Figure 8.**
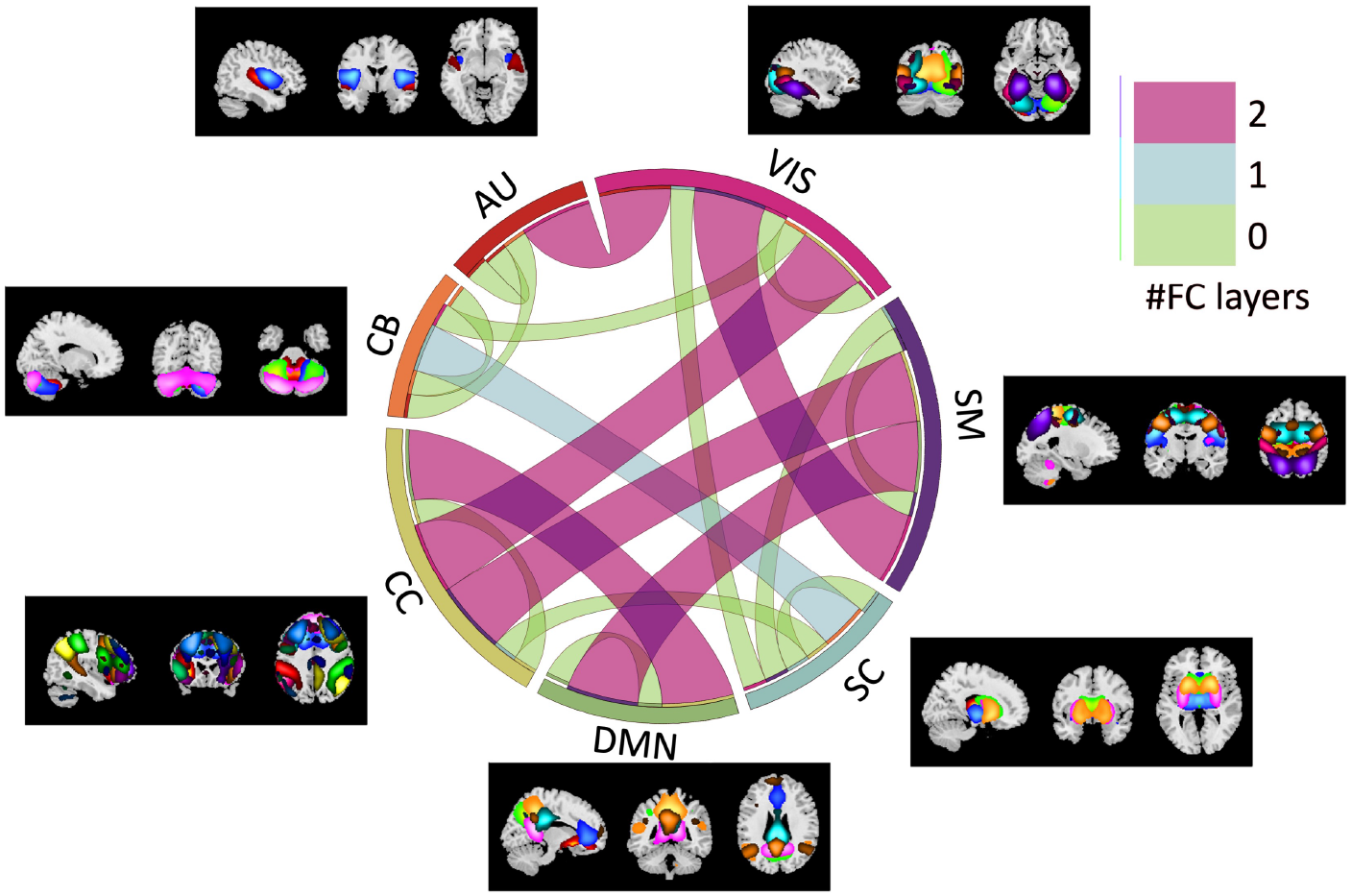
Connectogram showing SDIs requiring the same depth i.e., number of fully connected layers, in the TPE-optimized architectures for both COBRE and fBIRN datasets. The depth required by SDIs was the same for 19 out of 28 SDIs, indicating a common pattern across datasets in terms of certain SDIs requiring deeper models while others requiring shallower ones for a better prediction.

## 4 DISCUSSION

In this work we focus on developing a new approach for modeling differing values of variation and complexity with which information is encoded in the functional subdomain interactions (SDIs). We evaluate this approach in the context of a schizophrenia classification problem. The results show that allowing for differing subspace complexity can result in improved performance of the models (Figure 5), and enables us to identify meaningful differences in the modeling of different subdomains in the data, effectively using the resulting model complexity to determine whether their data contains more or less complex information about the prediction (Figure 8), given the model performance is at par with baseline frameworks. This trend of variation in the nature of information encoded in various functional subdomain interactions is significantly consistent (*p*-value = 2.09 ×10^−4^) across the independent datasets used in this study, our findings showed that 19 out of the 28 SDIs had the same depth in the optimized architectures for both the datasets.

Notably, it can be observed that the connections from the cognitive control (CC) to the visual (VIS) and sensorimotor (SM) subdomains require deeper models for both datasets (Figure 8(a)), indicating more complex models are needed to capture links between higher-order cognition (CC) and lower-order sensory areas (VIS, SM). Connectivity of components from SM, VIS and CC subdomains is well known to be affected in schizophrenia (Kaufmann et al., 2015; Javitt and Freedman, 2015). In fact, differences have been observed for both the focal (within SM and VIS components) as well as distal (between SM/VIS and CC components) connectivity in schizophrenia (Kaufmann et al., 2015; Javitt and Freedman, 2015;Butler and Javitt, 2005; Gaebler et al., 2015). The observation of auditory (AUD) subdomain to VIS subdomain connections requiring deeper models is interesting given previous work highlighting the disruptions in these areas in schizophrenia (Lynall et al., 2010; Calhoun et al., 2009; Gallinat et al., 2002;Rotarska-Jagiela et al., 2010; Yu et al., 2012) are connected in a complex manner and are implicated in auditory hallucinations and visual saccades, both of which are disrupted in schizophrenia, respectively. Changes in connectivity between the default mode network (DMN) and cognitive control (CC) areas are also well known, especially between the precuneus and the prefrontal cortex (Wolf et al., 2011). Interestingly, all SDIs involving the functional connectivity between components from the same network (i.e., self-connections in Figure 8) require shallower models, indicating that connections within a given network may encode less complex information compared to SDIs with inter-network connections, even if the former (intra-network connections) are known to be involved in schizophrenia (Garrity et al., 2007).

The TPE-optimized architectures perform significantly better than the corresponding architectures that do not allow for any variation in the depth required for various SDIs (UNIF0, UNIF1). Moreover, we also show by comparison with baseline methods that the multi-branched architecture also learns features that are consistent in terms of their discriminative power. Unlike the standard deep learning architectures such as MLP, it is possible to track feature importance in linear classifiers. However, the framework presented in this work is not only consistent with linear models in terms of feature importance, but also additionally allows for analyzing the variation in the nature of information in feature subdomains. Results from our work further emphasize the importance of questioning the assumption that various features from data as complex as neuroimaging data would require architectures with the same complexity irrespective of the subdomains of the brain. While it is apparent that certain subdomains contain more complex information for a given application, we provide an interpretable framework that analyzes as well exploits this variability for better understanding and classifying a given condition, in this case schizophrenia. A similar analysis could be extended to other brain disorders and diseases in future work, providing biomarkers for these conditions in terms of complexity of information in the subdomains of the brain.

Future work should also be focused on extending this analysis to study more parameter optimization methods as well as introducing more types of flexibility in the frameworks. Approaches to automatically determine the model structure/depth and including additional measures of model complexity would be quite powerful and is not well studied. Furthermore, developing flexible architectures like these does not only give insights towards identifying relationships between brain subdomains of the same modality, but it can also be extended to create adaptable yet rigorous frameworks that exploit complex interrelationships from subdomains in a multimodal setting.

## ACKNOWLEDGEMENTS

Research reported in this publication was supported by National Institute of Mental Health (HHS - NIH) of the National Institutes of Health under award numbers R01MH118695, RF1AG063153, and R01EB020407.

